# Life-long Dietary Restrictions have Negligible or Damaging Effects on Late-life Cognitive Performance: A Key Role for Genetics in Outcomes

**DOI:** 10.1101/2022.04.09.487742

**Authors:** Andrew R. Ouellette, Niran Hadad, Andrew Deighan, Laura Robinson, Kristen O’Connell, Adam Freund, Gary A. Churchill, Catherine C. Kaczorowski

## Abstract

Several studies report that caloric restriction (CR) or intermittent fasting (IF) can improve cognition, while others report limited or no cognitive benefits. Here, we compare the effects of 20% CR, 40% CR, 1-day IF, and 2-day IF feeding paradigms to ad libitum controls (AL) on Y-maze working memory and contextual fear memory (CFM) in a large population of Diversity Outbred mice that model the genetic diversity of humans. While CR and IF interventions improve lifespan, we observed no enhancement of working memory or CFM in mice on these feeding paradigms, and report 40% CR to be damaging in the context of long-term memory. Using Quantitative Trait Loci mapping, we identified the gene *Slc16a7* to be associated with late-life long-term memory outcomes in mice on lifespan promoting feeding paradigms. Limited utility of dieting and fasting on memory in mice that recapitulate genetic diversity in the human population highlights the need for anti-aging therapeutics that promote cognitive function, with a neuronal monocarboxylate transporter encoded by *Slc16a7* highlighted as novel target.

## Introduction

Aging remains the greatest risk factor for the onset of dementia in the human population (Swerdlow, 2007). Given the current and projected increase of lifespan in the population, the risk of developing age-related disorders, such as dementia, increases in parallel (Christensen et al., 2009; Lunenfeld and Stratton, 2013). As such, the demand for interventions of cognitive decline are higher than ever. Dietary interventions, such as caloric restriction (**CR**) and intermittent fasting (**IF**), have been proposed as potentially actionable treatments for age-related cognitive disorders in both humans and animal models (Halagappa et al., 2007; Kishi et al., 2015; Kuhla et al., 2013; Leclerc et al., 2020; Witte et al., 2009). However, results from these studies are often conflicting; many studies report that CR improves cognition while others report no effect (Burger et al., 2010; Dal-Pan et al., 2011; Harder-Lauridsen et al., 2017; Martin et al., 2007; Pifferi et al., 2018; Scott et al., 2014). We hypothesize that the divergence of these results may, in part, be explained by a lack of genetic diversity in many animal studies which often use a single or a few inbred strains, in addition to the inability to control environmental factors (i.e. access to healthcare, education, income, population structure etc.) in human studies that often disrupts expected outcomes (Watts et al., 2011).

The Diversity Outbred (**DO**) mouse population offers the opportunity to study the effects of CR and IF on cognitive aging within a genetically diverse population while still controlling environmental factors in a laboratory setting. The DO population is derived from 8 parental inbred lines, segregating for over 40 million single nucleotide polymorphisms that provide a great magnitude of genetic variation (Churchill, Gary A. et al., 2012). In this study, a population of 960 DO mice were separated into 5 feeding paradigms: *ad libitum* (**AL**) control, 20% or 40% CR, and 1 or 2-day IF. To assess the effect of aging, diet, and possible age x diet interactions on working memory, mice underwent longitudinal Y-maze working memory tests at 10 and 22-months of age (Fig 1A). Additionally, we assessed acquisition and recall of hippocampal-dependent spatial episodic memory using contextual fear conditioning at 24 months of age. Previous studies have shown effects of long-term CR on survival in mice can be observed by 22-24 months of age and is the approximate median lifespan of the DO population(Cameron et al., 2012; Mitchell, Sarah J. et al., 2019; Sun et al., 2013; Weindruch and Sohal, 1997), we therefore wanted to observe any potential changes in cognition at this age. Our results highlight the need for further mechanistic understanding of cognitive longevity to identify novel therapeutics.

**Figure 1.**
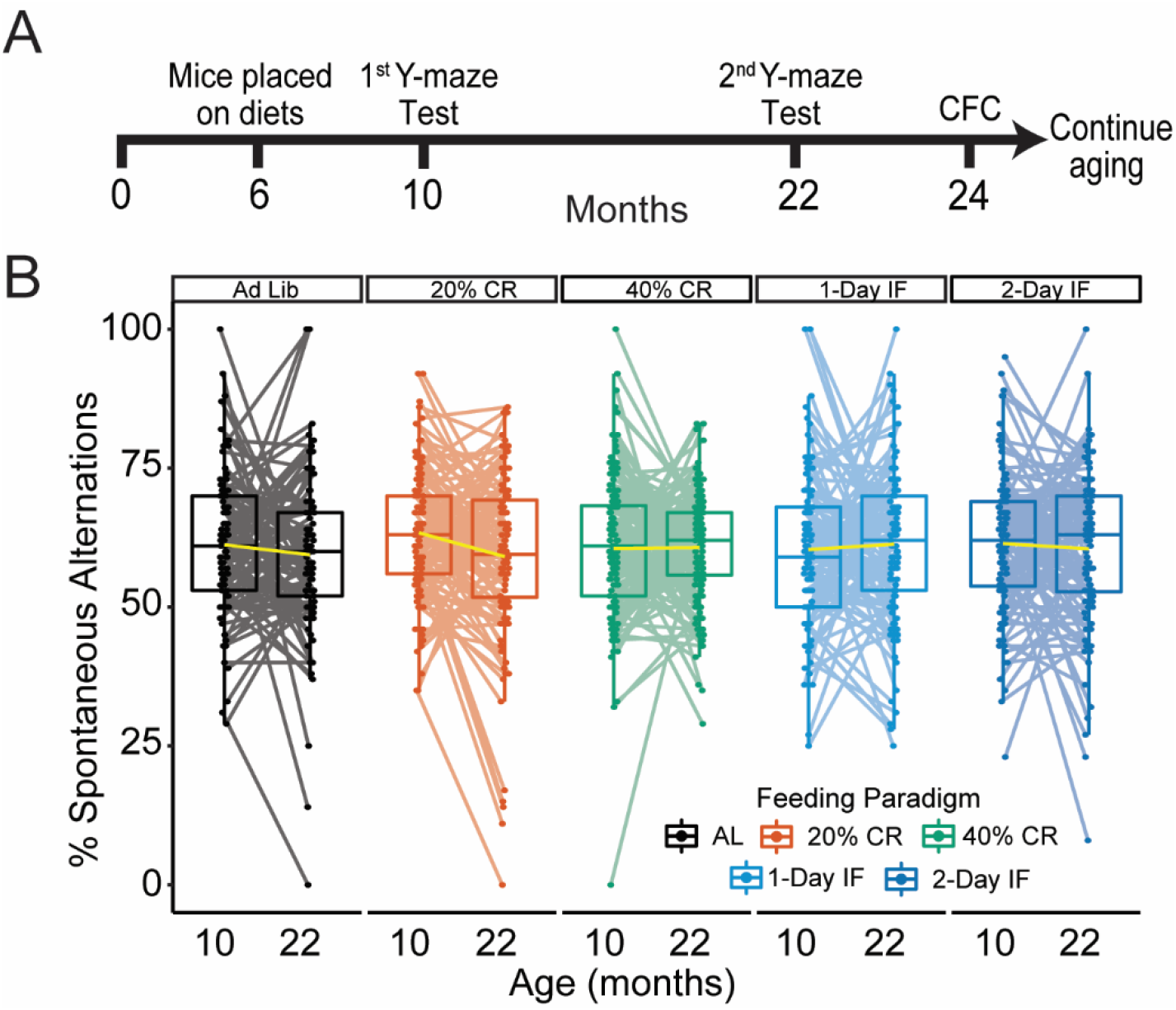
Caloric restriction and intermittent fasting paradigms have no effect on working memory in the Diversity Outbred Mouse population. (**A**) Overview of the experimental timeline; mice were longitudinally tested at 10 and 22 months on the Y-maze working memory task. After fear conditioning at 22 months mice were aged out to their maximum lifespan. (**B**) We observed no significant effects of diet on age related decline of working memory measured by % spontaneous alternations in the Y-maze working memory task from 10 months of age to 22. Each dot represents an individual mouse, yellow lines denote line of best fit between ages. Significance testing was performed using 2-way repeated measures ANOVA.

## Methods and Materials

### Animals

Female DO mice, obtained from the Jackson Laboratory, were used in this study (n = 960, J:DO, JAX stock number 009376, generations 24-28) previously described in (Churchill, G. A. et al., 2012) as part of a longitudinal maximum lifespan study. Mice were housed in polycarbonate cages on ventilated racks providing 99.997% HEPA filtered air to each cage in a climate-controlled room (ambient air temperature of 72°F) under a standard 12:12 light-dark cycle (lights on at 0600 h). Mice were housed together in groups of 8 and provided with wood blocks and plastic tubes for enrichment. At 6 months of age mice were placed into one of the following 5 diet cohorts (n = 192 in each cohort). 1) *Ad libitum* 2) 20% Caloric restriction (2.75 g/mouse/day), 3) 40% Caloric Restriction (2.06g/mouse/day), 4) 1 Day Fast (food removed Wednesday 15:00 and given Thursday 15:00), 5) 2 Day Fast (food removed Wednesday 15:00 and given Friday 15:00). Caloric restricted mice were provided food based on the number of mice in a given pen, mice in these pens were provided food at the same time. All mice were provided with 6% fat extruded grain (5K0G chow, LabDiet, St. Louis, MO).

### Genotyping

Tail tip samples were collected DNA extracted with the DNeasy Blood and Tissue Kit (Qiagen) from 954 animals (6 mice died before genotyping). Extracted DNA was genotyped on a 143,259-probe GigaMUGA array from the Illumina Infinium II platform (Morgan et al., 2016) by NeoGen Corp. Genotype quality was evaluated with the R qtl2 package (Broman et al., 2019). All the raw genotype data was processed with a corrected physical map of the GigaMUGA array probes. After processing the genotype dataset contained 110,807 markers.

### Y-maze Working memory Task

At 10 and again at 22 months of age, all mice were allowed to freely explore a Y-maze apparatus with three equally-sized arms for 8 minutes: arm length of 30 cm, arm lane width of 5-6 cm, wall height of 12-18 cm. External visual cues were removed from the apparatus with a curtain which surrounded the apparatus. Working memory was measured by % spontaneous alternations (Number of spontaneous alternations / Total arm Entries) between Y-maze arms. Recorded videos were analyzed in ANY-maze behavioral tracking software (Stoelting Co., IL, United States). To limit the number of false positive arm entries we counted an arm entry when 99% of the mouse’s body (excluding tail) crossed into a new arm. Mice that did not make at least 6 total arm entries (the minimum number needed for at least 4 spontaneous alternations) were excluded from Y-maze data analysis.

### Contextual Fear Conditioning

At 24 months of age all mice underwent Contextual Fear Conditioning (CFC) to assess hippocampal-dependent short and long-term memory (Neuner et al., 2015). On the first day of training, mice were placed in a training chamber and four foot-shocks (0.9 mA, 1 s) were delivered after a 150 second baseline period. Four post-shock intervals were defined as the 40 s following the end of each foot shock and the percentage of time spent freezing during each interval was determined using FreezeFrame software (Actimetrics Inc., IL, United States). The percentage of time spent freezing following the final shock was used as a measure of contextual fear acquisition across the panel. Twenty-four hours after training, mice were placed back into the training chamber and the percentage of time spent freezing throughout the entire 10-min test was measured as an index of contextual fear memory; no shocks were delivered during the testing session. Mice were habituated in the testing room for 1 hour before testing/training on both day 1 and 2. Mice were fear conditioned on Monday/Tuesday to avoid testing during fasting periods.

### Data Analysis

Statistical analysis was performed using R 4.1.1. <u>Survival</u>: Mice that did not die from age-related causes (i.e. Missing or accidental death) were excluded from survival analysis. We fitted a logistics regression model with survival as a response variable and feeding paradigm as a fixed effect. This model accounts for clustering of mice into pens by using a nested random effect of pen within DO generation (given that any generation contains only a specific set of pens). P-values from the mixed model were corrected for multiple testing using Benjamini-Hochberg correction. <u>Y-maze</u>: To test for differences in age related decline of working memory from 10mo to 22mo between diet cohorts we utilized a 2-way repeated measures ANOVA; only mice that were in both age groups were analyzed. <u>CFA</u>: Slope acquisition derived from the coefficient of a linear model with the 4 post-shock values fitted against values ranging from 1-4. To test for differences in contextual fear conditioning data between diets we utilized linear mixed modeling with the Lme4 R package (Bates et al., 2015). <u>CFM</u>: To test for a significant effect of diet on CFM we used linear mixed modeling with Lme4. In this model diet was treated as a fixed effect, test date as a random effect, and pen and generation as nested random effects. Post hoc p-values were derived from Tukey HSD testing and corrected for multiple comparisons among the 5 diet cohorts using Holms correction for both CFA and CFM. <u>Heritability</u>: Heritability of cognitive phenotypes was calculated via the qtl2 “est_herit” function (Version 0.28) (Broman et al., 2019) by fitting a mixed-effects model controlling for feeding paradigm as a fixed effect and utilizing a genetic kinship matrix as a random effect.

### QTL Mapping

We performed QTL mapping and SNP association mapping with R using the qtl2 package (Version 0.28) (Broman et al., 2019). Log of the odds ratio (LOD) was derived from a linear mixed model and reported mapping statistic. The significance thresholds for QTL were calculated with the “scan1_perm” function in qtl2 with 1000 permutations. For each permutation the genotype probabilities for each mouse are randomly assigned to different covariate and the maximum LOD score for the proceeding genome scan is recorded. To determine statistical significance of a QTL peak, a genome-wide p-value of 0.05 (95^th^ percentile of recorded maximum LOD scores) from permutation testing was used. In genome scans, permutations and SNP association, we accounted for genetic similarity between mice using a kinship matrix with the leave-one-chromosome-out (LOCO) method. The founder allelic effect was identified using a regression of the phenotype on the founder genotype probabilities at each locus. For all CFM mapping, date of testing was used as an additive covariate to control for observed batch effects. To test for significant QTL across the entire DO population we used the 5 feeding paradigms as an interactive covariate. To test for significant QTL in mice on lifespan promoting and long-term memory preserving diets the DO population was separated to mice on 20%CR, 1-day IF and 2-day IF. These feeding paradigms were treated as a single covariate; therefore, no interactive mapping was used.

## Results

### Diversity Outbred mice do not experience age-related working memory decline in a free exploring Y-maze task

The loss of working memory in aged humans and mice is well characterized in previous studies (Erickson and Barnes, 2003; Kirova et al., 2015; van Geldorp et al., 2015). However, the effect of CR and IF on working memory remains a point of contention in the literature. Here, we assessed working memory in young (10 month) and mid-aged (22 month) DO mice on AL, calorically restricted and intermittent fasted paradigms by measuring percent spontaneous alternation (**%SA**) in the Y-maze working memory task. We found that while DO mice on the control AL paradigm exhibited a general trend towards an age-related decrease in %SA from 10 and 22 months this effect was not statistically significant (p>0.05), nor was this trend affected by CR or IF interventions in adult (10mo) or aged (22mo) DO mice (Fig. 1B). It is important to note that spatial cues were not provided to mice during free exploration of the Y-maze. In this configuration, the Y-maze task is not hippocampal dependent and, therefore, not sensitive to age-related cognitive decline (Albani et al., 2014). As such, we are unable to determine if the absence of age-related decline is imparted by the lack of either spatial reference in the apparatus or true cognitive deficits (Lennartz, 2008).

### Intermittent fasting confers no Contextual Fear Memory benefits while severe caloric restriction is damaging in aged Diversity Outbred mice

Age-related decline in acquisition, consolidation and recall of spatial and contextual memories are known to be associated with hippocampal dysfunction in both animal models and humans (Park et al., 1996; Wimmer et al., 2012). To assess the effectiveness of CR and IF interventions in the preservation of performance on hippocampal-dependent short-term and long-term memory relative to the AL controls in the DO mice, we performed contextual fear conditioning at 24 months. Acquisition of short-term contextual fear memory was measured as the change in percent time spent freezing of the mice after each of 4 foot shocks on day 1 of training (Fig. 2A, top), previously characterized as a measurement of memory acquisition (Neuner et al., 2019). We found that mice acquire short-term contextual fear acquisition (**CFA**, measured by slope acquisition and % freezing during post-shock 4) on day 1 of training comparably regardless of feeding paradigm, suggesting that neither long-term CR nor IF have a profound effect on short-term memory performance relative to AL (Fig. 2B, Fig. S1).

**Figure 2.**
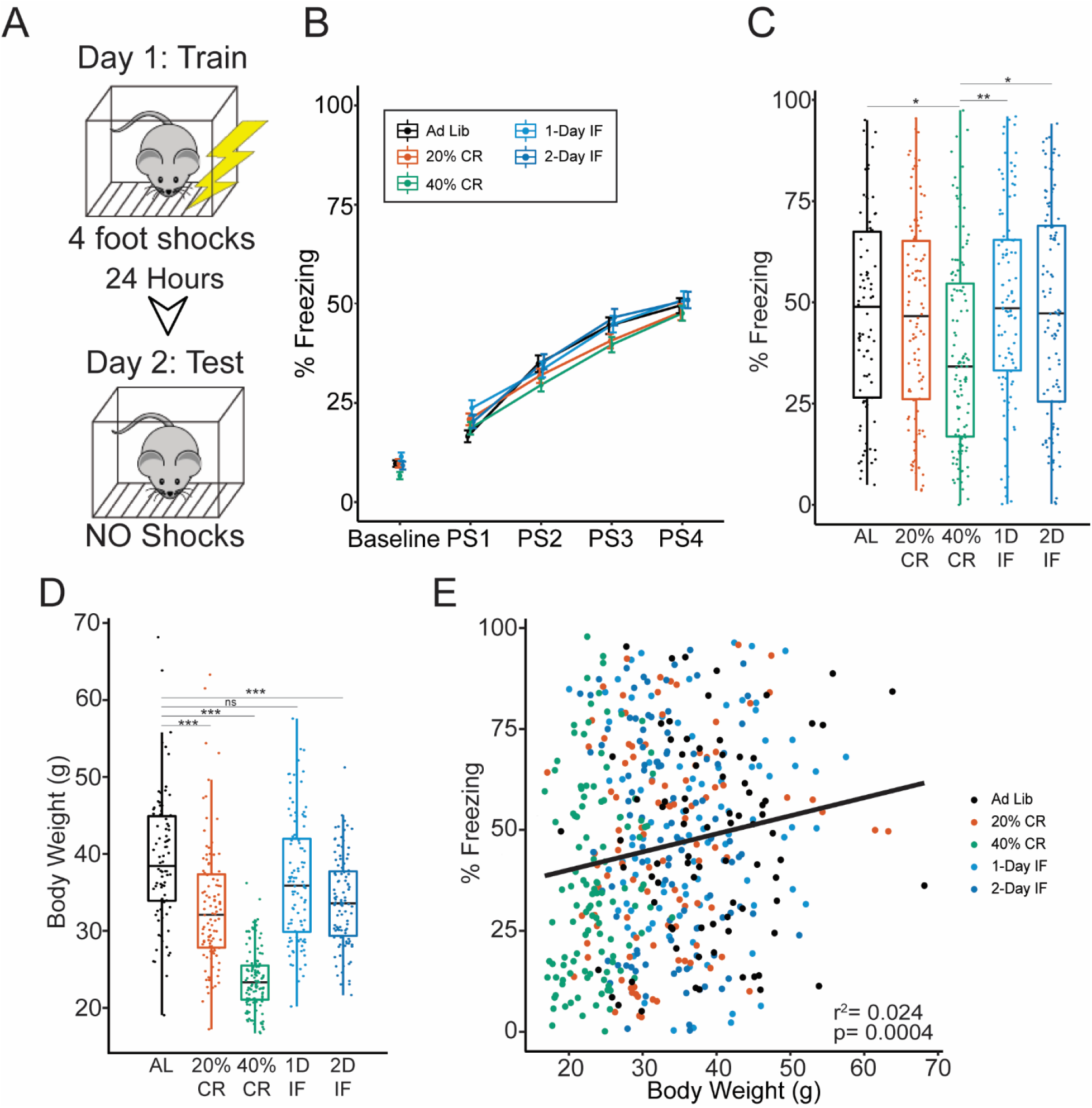
Contextual Fear Memory is impaired in diversity outbred mice on 40% caloric restriction compared to intermittent fasting and Ad Lib control groups. (**A**) CFC testing schematic; on day 1 mice were given 4 foot-shocks. Twenty-four hours later mice were placed in the same context without any shocks (**B**) % Freezing on day 1 of training before receiving shocks (baseline) and during each post-shock (PS) interval. Acquisition of CFC was comparable across all feeding paradigms. Data are expressed as mean ± SEM, significance was determined using 1-way repeated measures ANOVA. (**C**) Contextual Fear Memory (CFM) measured by total percent freezing of mice on day 2. Significant effect of diet on % Freezing was determined with a linear mixed model with test batch treated as a random effect. (**D**) Boxplot of body weights at 24 months of age across all feeding paradigms. A Post-hoc Tukey test was used to determine significant differences between diet groups; p-values were corrected for multiple comparisons using Holm correction (**E**) Scatter plot correlating CFM percent freezing and 24mo body weight across the entire DO population. ^ns^p>0.05, *p<0.05, **p<0.01 ***p<0.001, n = 502. Boxplots encompass the 25th to 75th percentile with whiskers indicating 10th and 90th percentiles, median lines are indicated within each box.

To assess the effects of CR and IF on long-term contextual fear memory (**CFM**), we measured the total percent freezing of the mice during the day-2 testing period (Fig. 2A, bottom). To test for a main effect of feeding paradigm on CFM performance we used a linear mixed model that accounts for testing batch effects as a random effect in addition to nested pen density within different generations (F(4,502) = 4.451, p < 0.01, Fig. 2C, Fig. S2). Post hoc analysis revealed no significant differences between AL controls and feeding interventions, except for the 40% CR cohort, which exhibited impaired CFM (Fig. 2C) (Tukey HSD, p = 0.03, 95% C.I. = AL [40.3, 54.9], 40%CR [30.3, 43.9]). We also observed that CFM in 40% CR was still significantly impaired compared to both IF interventions (Tukey HSD, p < 0.02, 95% C.I. = 40%CR [30.3, 43.9], 1D-IF%CR [41.9, 55.8], 2D-IF [40.6, 54.5], Figure 2C). These results suggest that, while CR and IF interventions reported herein replicate lifespan extension using percent survival at 24 months as a proxy measure (Table 1, (Chi-Square=17.82 on 4 df, p=.001)), CR and IF do not promote late-life cognitive performance compared to AL feeding, and in the case of 40% CR, is likely damaging compared to moderate 20% CR and IF interventions.

**Table 1.** Caloric restriction and intermittent fasting paradigms increase percent survival at 24 months of age. Proportion of mice that survived until 22 months within each diet cohort. Mice that died from non-natural causes (i.e missing animals, mishandling and non-endpoint euthanasia) were excluded from survival analysis. We observed that all diet interventions conferred increased survival compared to ad lib controls, 40% CR confers the greatest % survival [χ^2^ (4, n = 861) = 17.4, p = 0.002]. Post hoc testing was performed with multiple chi-squared tests comparing diet cohorts to ad lib controls, Holm multiple testing correction was used to correct multiple testing error. *p < 0.05, **p<0.01 ***p<0.001

| Feeding Paradigm | % Survived to 24mo | Total n included | p-value compared to Ad Lib |
| --- | --- | --- | --- |
| Ad lib | 46.3 | 175 | NA |
| 20% CR | 59.3 | 172 | * |
| 40% CR | 68.0 | 169 | *** |
| 1-day IF | 59.0 | 173 | * |
| 2-day IF | 60.5 | 172 | ** |

In any feeding study changes in and correlation with body weight also need to be considered. As expected, we observed a significant effect of feeding paradigm in body weight at the median 24-month timepoint (F(4,502) = 76.01, p < 0.0001, with 40% CR imparting the most severe reduction in body weight when compared to AL controls (Tukey HSD, p < 0.0001) Fig. 2D). While individual changes in body weight contribute little variance to CFM outcomes (2.4% variance explained), we detected a significant positive correlation between these two traits (Fig. 2E), suggesting that metabolic outcomes altered by diet are coupled with late-life cognition.

The variance in CFA/CFM performance in this DO population is much higher than inbred mouse strains given the same conditioning, which can attributed to the population’s enhanced genetic diversity (Bolivar et al., 2001; Tuttle et al., 2018). Similar distribution in phenotypes in previous DO studies and have pointed to the significant impact of heritability (**h^2^**) on phenotypic outcomes (Keenan et al., 2020). To determine whether genetic variation can explain the high degree of variance in CFA/CFM, we calculated heritability for these traits using a linear mixed modeling approach with the R qtl2 package (Broman et al., 2019). When accounting for variance between feeding paradigms, we calculate that 54.7% of the CFM variance can be explained by genetic diversity. This h^2^ estimate is much higher than Y-maze %SA providing further evidence that the lack of a WM phenotype was based in environmental factors, such as visual cues (Table 2). We propose this high heritability not only makes large CFM variance permissible, but also encouraging, given that we expect individual genetic background to influence late-life cognitive outcomes. We next sought to determine if specific regions in the genome were associated with late-life cognitive outcomes and if any specific genes or variant could provide actionable therapeutic targets.

**Table 2.**
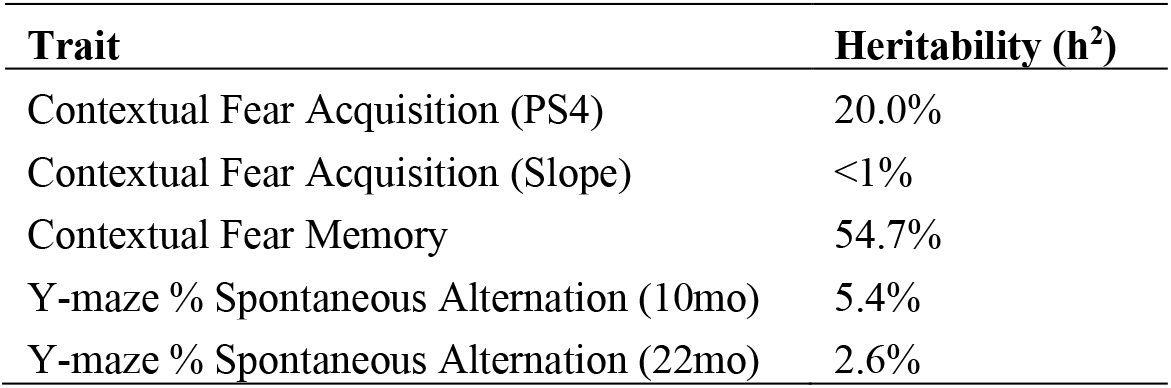
Heritability of cognitive phenotypes within the DO population. h^2^ is presented as a percent of the variation in a phenotype that can be explained by differences in genetic background across the DO population. A higher percentage is indicative of a greater percentage of variability explained by genetics

| **Trait** | **Heritability (h<sup>2</sup>)** |
| --- | --- |
| Contextual Fear Acquisition (PS4) | 20.0% |
| Contextual Fear Acquisition (Slope) | <1% |
| Contextual Fear Memory | 54.7% |
| Y-maze % Spontaneous Alternation (10mo) | 5.4% |
| Y-maze % Spontaneous Alternation (22mo) | 2.6% |

### Slc16a7 is associated with long-term memory outcomes in mice on feeding paradigms that promote lifespan and preserve cognitive function

While genetic background has been shown to modify cognitive declines in aged individuals, the exact genetic factors which underly these outcomes remain poorly understood. As such, we performed Quantitative Trait Loci (QTL) mapping across our DO population to identify loci that are associated with worsened or enhanced cognitive outcomes in the face of dietary interventions. Mapping of Y-maze working memory, CFA or CFM with feeding paradigm as an additive and/or interactive covariate yielded no significant QTL peaks (data not shown) across the genome. The lack of a significant QTL peak in the highly heritable CFM data can, in part, result from of high degrees of freedom within our interactive QTL mapping (comparisons between 5 diet groups). In order to reduce the degrees of freedom within our QTL mapping, while still testing for genetic associations with behavior in mice that were on lifespan promoting feeding paradigms which did not harm cognitive outcomes, we mapped only 20% CR, 1-day and 2-day IF cohorts as a single covariate. QTL mapping of late-life CFM outcomes in these mice revealed a significant peak near the end of chromosome 10 ranging from 124.2-125.3Mbp (alpha=0.05, Fig. 3A). Within this locus we identified several gene modules, a Riken gene and the protein coding gene ***Slc16a7*** (Solute carrier family 16 member 7) (Fig. 3B, top). Additionally, we identified 256 single nucleotide polymorphisms **(SNPs)** within this region associated with changes in cognitive outcomes and are found in 3 of the DO founder strains (AJ, NZO and WSB) (Fig. 3B Bottom). 4 of these SNPs are upstream of GM23777 and 2 downstream, the remaining 250 SNPs are classified as intergenic. Based on previous research investigating the role of *Slc16a7* in metabolism and cognitive performance (Lev-Vachnish et al., 2019; Lu et al., 2015; Netzahualcoyotzi and Pellerin, 2020; Yu et al., 2021), we prioritized this gene over identified gene modules. We next sought to determine if these SNPs were associated with enhanced or worsened late-life cognitive outcomes. Using a linear mixed model that accounts for kinship, we calculated QTL effects from each of the 8 DO founders along chromosome 10. We found that mice with AJ, NZO and WSB alleles in the QTL peak region had higher late life cognitive function than mice with B6 or NOD alleles (Fig. 3C). These results indicate that variants within this region are with enhanced late-life cognitive performance in individuals on lifespan and cognitive health span promoting dietary interventions.

**Figure 3.**
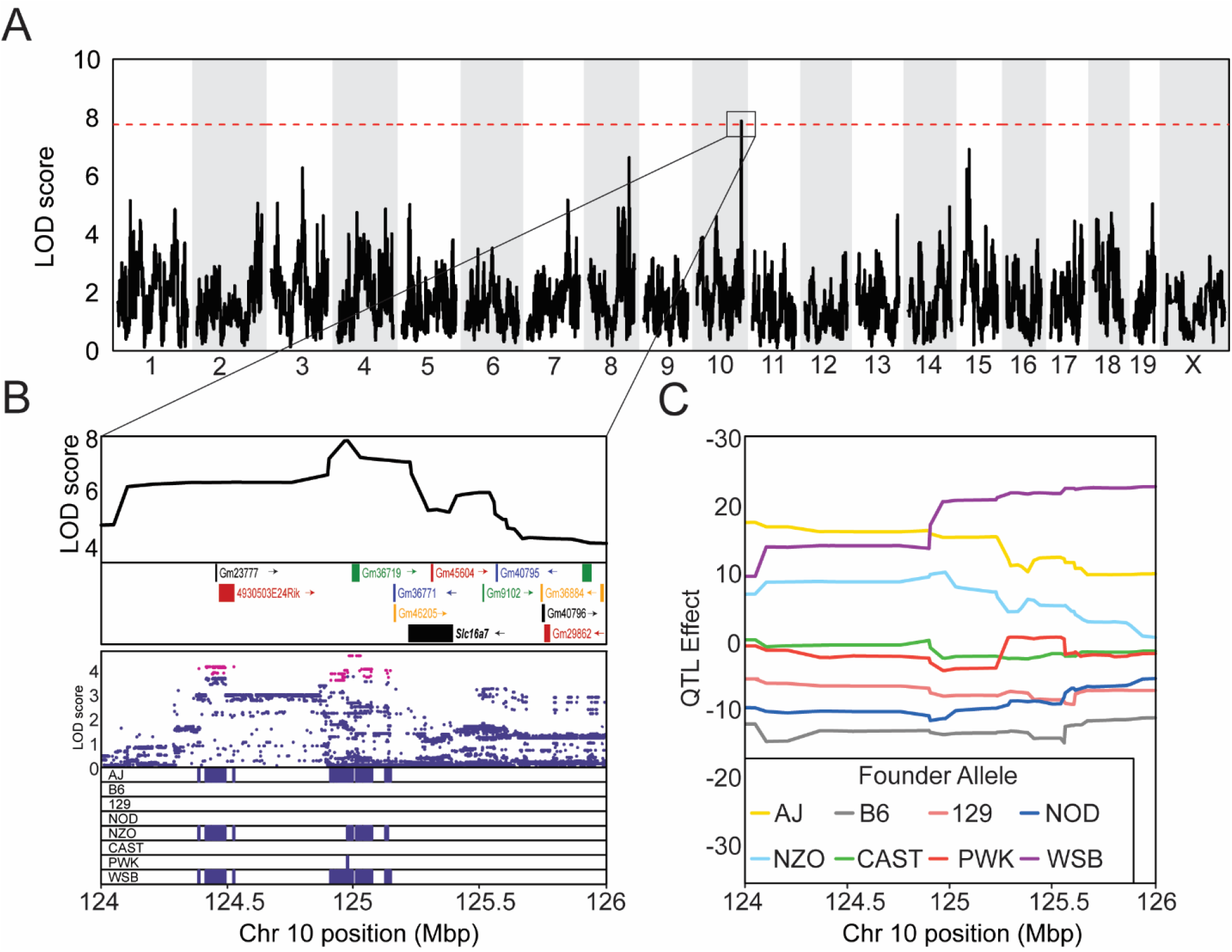
Slc16a7 is associated with variation long-term memory outcomes. (**A**) QTL map across the entire genome identifies a locus on chromosome 10 that significantly associates with changes in late-life CFM. Significance score (LOD) is represented on the y-axis. The dashed red represents significance threshold from permutation testing (1000 permutations, p-value = 0.05) (**B**) (Top) Area under the significant QTL peak showing genes and gene modules within this region. (Bottom) SNP marker associations of the eight DO founders labeled above. The x-axis shows the distribution along the chromosome in physical distance, SNPs colored in red fall within a LOD 1.5 drop of the peak marker. Minor allele frequency for SNPs significant SNPs is shown below. The x-axis is physical distance in Mb along the chromosome. (**C**) Haplotype effects of the 8 founders at the QTL region for CFM. The y-axis for the top panel is the effect coefficient

## Discussion

Heritability estimates for age-related cognitive decline, including the onset of dementias such as Alzheimer’s disease are in the range of 58-79% (Deary et al., 2012; Gatz et al., 2006; Harris and Deary, 2011; Ridge et al., 2013), suggesting that genetic background plays an integral role in regulating cognitive aging. Studies that use traditional inbred rodent models of aging fail to capture the impact of genetic diversity on cognitive traits, resulting in a lack of translatability to the genetically diverse human population. Additionally, many of these studies reporting cognitive improvement in CR treated cohorts report only marginal improvement in memory tasks that assess only a single cognitive domain (Dhurandhar et al., 2013; Parikh et al., 2016; Wahl et al., 2018; Wu, Aiguo et al., 2003). Using a large panel of genetically diverse mice allowed us to robustly evaluate the effects of feeding interventions on cognitive performance across genetic backgrounds, enhancing the likelihood that results will generalize across diverse populations and species. Moreover, our results suggest genetic background may be a key factor in conflicting findings in previous reports using model organisms of aging, given the complex nature of gene x diet interactions that are known to modify cognitive outcomes in genetically diverse models of dementia (Tucker-Drob et al., 2013).

With our reported gene mapping results, we highlight the utility of DO mice in investigating the interaction between genetic background and late-life cognitive outcomes. Mapping of late-life long-term memory performance successfully identified *Slc16a7* as a potential mediator of cognitive outcomes in individuals undergoing lifespan promoting CR and IF interventions. *Slc16a7*, which codes the lactate and pyruvate transporter protein **MCT2** (Monocarboxylate transporter 2), has previously been implicated age-related cognitive disfunction in rodents and humans via neuronal lactate transportation (Lev-Vachnish et al., 2019; Lu et al., 2015; Netzahualcoyotzi and Pellerin, 2020; Yu et al., 2021). It is known that lactate is an important energy source for de novo mRNA translation, a process critical for long-term memory consolidation (Descalzi et al., 2019), and that MCT2 functions in hippocampal neurons to achieve this demand for energy (Netzahualcoyotzi and Pellerin, 2020). We propose that genetic diversity, in conjunction with moderate caloric restriction and intermittent fasting, in the region of *Slc16a7* is a driver for late-life cognitive resilience. If there are intergenic variants that are affecting the expression of MCT2 in the DO population, it stands to reason that these changes in individual expression may impart observed cognitive enhancement or detriments. Future studies will investigate the role of these variants in altering the expression of MCT2 and the modification of cognitive outcomes, and whether/if they are altering the function of MCT2 proteins or their level of expression in hippocampal neurons. Given the existing evidence of *Slc16a7* as a mediator of cognitive outcome, we propose it to be the gene of interest from our QTL mapping rather than identified gene modules. However, because significant variants that associate with enhanced cognitive outcomes are found within and around other gene modules and Riken genes, further investigation into GM36719, GM23777 and 4930503E24Rik is warranted and may yield new therapeutic targets and knowledge of late-life cognitive outcomes. Identification of *Slc16a7* in conjunction with the high heritability of late-life contextual fear memory demonstrates that the DO mouse population can effectively model the aging process across a genetically diverse population and shows that this process can be linked to previously confirmed genetic mechanisms.

While CR and IF are highly replicable and well established interventions of health-span and lifespan across multiple species ranging from worms to humans (Catterson et al., 2018; Lakowski and Hekimi, 1998; Mattison et al., 2017; Mitchell, S. J. et al., 2019; Most et al., 2017; Rogina and Helfand, 2004) and in our own data, their efficacy to promote late-life cognitive performance has been mixed; the reason for this remains largely unexplored. There are many factors that may interact with caloric restriction and modify its efficacy as an intervention of cognitive decline, including genetic background and sex. Previous studies have shown that caloric restriction is more effective at promoting lifespan in certain inbred mouse strains, while less effective in others (Liao et al., 2013; Liao et al., 2010; Mitchell et al., 2016; Wilkie et al., 2020), suggesting that certain individuals may be predisposed to effectively utilizing caloric restriction strategies. Our work has shown that these methods are not effective at promoting late-life cognitive function on a population level. Additionally, previous studies have shown that sex modifies the effect of dietary restrictions on maximum lifespan and cognition (Kane et al., 2018; Mitchell et al., 2016; Wu, A. et al., 2003; Zajitschek et al., 2013). It is therefore important to consider gene x sex interactions when evaluating the efficacy of caloric restrictions on both lifespan and cognitive aging. Due to animal health and attrition concerns stemming from previously observed DO male aggression, only females were used in this study. Future work will need to include both males and females in order to determine gene x sex interactions that may affect the efficacy of dietary restrictions.

We also need to consider the role of survival bias in the interpretation of our results given that there is an increase of survival in the 40% CR compared to ad lib controls. The possibility that the genetic factors potentially decreasing survival in the ad libitum group also enhanced their cognitive abilities is a valid consideration. However, since DO mice were randomized across feeding cohorts, we expect an equal distribution of genotypes within each cohort. Furthermore, if a relationship between survival and cognition was expected, and may be a source for survival bias, we would 1. Expect to see it in other pro-longevity paradigms (1/2 day IF and 20% CR) and 2. Pro-longevity paradigm would result in improved cognitive outcomes. Therefore, if there was a selection bias against late-life poor cognitive performers in the ad lib cohort, we would expect to see a relatively more of these mice in enhanced survival cohorts and a reduction in cognitive outcomes in the 20% CR, 1-day and 2day IF cohorts; this is not the case.

## Conclusion

DO mice have been shown as an effective tool in targeting genetic mechanisms of aging that reliably translate into humans (Church et al., 2015; French et al., 2015; Ouellette et al., 2020; Recla et al., 2014; Tuttle et al., 2018). We, therefore, highlight the need to investigate alternative therapeutics of not only lifespan, but also cognitive health-span, given that CR and IF are proving to fall short of desired outcomes. In order to develop more effective therapeutics, we need to develop a better understanding of the genetic mechanisms of cognitive longevity; our future work will continue utilizing DO mice to do just this. By harnessing the complex genetic backgrounds of DO mice, we will investigate how genetic mechanisms regulate not only cognitive outcomes, but also molecular profiles (RNA expression and DNA methylation) and dendritic spine morphology that we and others showed associates with cognitive resilience in normal and AD aging (Kasai et al., 2010; Walker and Herskowitz). These studies will provide insights into new genetic targets for potential therapeutics to rescue cognitive deficits and dementia.

## Author contributions

Conceptualization: CCK, GC, AF, AO; Methodology: AO, CCK, NH, GC, AD; Investigation: AO, CCK, GC, AF; Visualization: AO, CCK, NH, KO, AD; Funding acquisition: CCK, GC, AF; Project administration: CCK, GC, AF, LR; Supervision: CCK, GC, AF; Writing – original draft: AO, CCK, NH; Writing – review & editing: AO, CCK, NH, KO, GC.

## Data and materials availability

Data used for analyses and scripts are available at request

## Abbreviations

CR: Caloric Restriction
IF: Intermittent Fasting
DO: Diversity Outbred
%SA: Percent spontaneous Alternations
CFM: Contextual Fear Memory
CFA: Contextual Fear Acquisition
LOD: Log of Odds
ANOVA: Analysis of Variance
SEM: Standard Error of Mean

## References

Albani, S.H., McHail, D.G., Dumas, T.C., 2014. Developmental studies of the hippocampus and hippocampal-dependent behaviors: insights from interdisciplinary studies and tips for new investigators. Neurosci Biobehav Rev 43, 183–190.

Bates, D., Mächler, M., Bolker, B., Walker, S., 2015. Fitting Linear Mixed-Effects Models Using lme4. Journal of Statistical Software; Vol 1, Issue 1 (2015).

Bolivar, V.J., Pooler, O., Flaherty, L., 2001. Inbred strain variation in contextual and cued fear conditioning behavior. Mammalian Genome 12(8), 651–656.

Broman, K.W., Gatti, D.M., Simecek, P., Furlotte, N.A., Prins, P., Sen, Ś., Yandell, B.S., Churchill, G.A., 2019. R/qtl2: Software for Mapping Quantitative Trait Loci with High-Dimensional Data and Multiparent Populations. Genetics 211(2), 495–502.

Burger, J.M., Buechel, S.D., Kawecki, T.J., 2010. Dietary restriction affects lifespan but not cognitive aging in Drosophila melanogaster. Aging cell 9(3), 327–335.

Cameron, K.M., Miwa, S., Walker, C., von Zglinicki, T., 2012. Male mice retain a metabolic memory of improved glucose tolerance induced during adult onset, short-term dietary restriction. Longevity & Healthspan 1(1), 3.

Catterson, J.H., Khericha, M., Dyson, M.C., Vincent, A.J., Callard, R., Haveron, S.M., Rajasingam, A., Ahmad, M., Partridge, L., 2018. Short-Term, Intermittent Fasting Induces Long-Lasting Gut Health and TOR-Independent Lifespan Extension. Current Biology 28(11), 1714–1724.e1714.

Christensen, K., Doblhammer, G., Rau, R., Vaupel, J.W., 2009. Ageing populations: the challenges ahead. Lancet 374(9696), 1196–1208.

Church, R.J., Gatti, D.M., Urban, T.J., Long, N., Yang, X., Shi, Q., Eaddy, J.S., Mosedale, M., Ballard, S., Churchill, G.A., Navarro, V., Watkins, P.B., Threadgill, D.W., Harrill, A.H., 2015. Sensitivity to hepatotoxicity due to epigallocatechin gallate is affected by genetic background in diversity outbred mice. Food and Chemical Toxicology 76, 19–26.

Churchill, G.A., Gatti, D.M., Munger, S.C., Svenson, K.L., 2012. The diversity outbred mouse population. Mammalian Genome 23(9), 713–718.

Churchill, G.A., Gatti, D.M., Munger, S.C., Svenson, K.L., 2012. The Diversity Outbred mouse population. Mammalian genome: official journal of the International Mammalian Genome Society 23(9-10), 713–718.

Dal-Pan, A., Pifferi, F., Marchal, J., Picq, J.L., Aujard, F., 2011. Cognitive performances are selectively enhanced during chronic caloric restriction or resveratrol supplementation in a primate. PloS one 6(1), e16581.

Deary, I.J., Yang, J., Davies, G., Harris, S.E., Tenesa, A., Liewald, D., Luciano, M., Lopez, L.M., Gow, A.J., Corley, J., Redmond, P., Fox, H.C., Rowe, S.J., Haggarty, P., McNeill, G., Goddard, M.E., Porteous, D.J., Whalley, L.J., Starr, J.M., Visscher, P.M., 2012. Genetic contributions to stability and change in intelligence from childhood to old age. Nature 482(7384), 212–215.

Descalzi, G., Gao, V., Steinman, M.Q., Suzuki, A., Alberini, C.M., 2019. Lactate from astrocytes fuels learning-induced mRNA translation in excitatory and inhibitory neurons. Communications biology 2(1), 247.

Dhurandhar, E.J., Allison, D.B., van Groen, T., Kadish, I., 2013. Hunger in the Absence of Caloric Restriction Improves Cognition and Attenuates Alzheimer’s Disease Pathology in a Mouse Model. PloS one 8(4), e60437.

Erickson, C.A., Barnes, C.A., 2003. The neurobiology of memory changes in normal aging. Experimental gerontology 38(1-2), 61–69.

French, J.E., Gatti, D.M., Morgan, D.L., Kissling, G.E., Shockley, K.R., Knudsen, G.A., Shepard, K.G., Price, H.C., King, D., Witt, K.L., Pedersen, L.C., Munger, S.C., Svenson, K.L., Churchill, G.A., 2015. Diversity Outbred Mice Identify Population-Based Exposure Thresholds and Genetic Factors that Influence Benzene-Induced Genotoxicity. Environmental Health Perspectives 123(3), 237–245.

Gatz, M., Reynolds, C.A., Fratiglioni, L., Johansson, B., Mortimer, J.A., Berg, S., Fiske, A., Pedersen, N.L., 2006. Role of Genes and Environments for Explaining Alzheimer Disease. Archives of General Psychiatry 63(2), 168–174.

Halagappa, V.K., Guo, Z., Pearson, M., Matsuoka, Y., Cutler, R.G., Laferla, F.M., Mattson, M.P., 2007. Intermittent fasting and caloric restriction ameliorate age-related behavioral deficits in the triple-transgenic mouse model of Alzheimer’s disease. Neurobiology of disease 26(1), 212–220.

Harder-Lauridsen, N.M., Nielsen, S.T., Mann, S.P., Lyngbæk, M.P., Benatti, F.B., Langkilde, A.R., Law, I., Wedell-Neergaard, A.S., Thomsen, C., Møller, K., Karstoft, K., Pedersen, B.K., Krogh-Madsen, R., 2017. The effect of alternate-day caloric restriction on the metabolic consequences of 8 days of bed rest in healthy lean men: a randomized trial. Journal of applied physiology (Bethesda, Md.: 1985) 122(2), 230–241.

Harris, S.E., Deary, I.J., 2011. The genetics of cognitive ability and cognitive ageing in healthy older people. Trends in cognitive sciences 15(9), 388–394.

Kane, A.E., Sinclair, D.A., Mitchell, J.R., Mitchell, S.J., 2018. Sex differences in the response to dietary restriction in rodents. Curr Opin Physiol 6, 28–34.

Kasai, H., Fukuda, M., Watanabe, S., Hayashi-Takagi, A., Noguchi, J., 2010. Structural dynamics of dendritic spines in memory and cognition. Trends in Neurosciences 33(3), 121–129.

Keenan, B.T., Galante, R.J., Lian, J., Simecek, P., Gatti, D.M., Zhang, L., Lim, D.C., Svenson, K.L., Churchill, G.A., Pack, A.I., 2020. High-throughput sleep phenotyping produces robust and heritable traits in Diversity Outbred mice and their founder strains. Sleep 43(5).

Kirova, A.M., Bays, R.B., Lagalwar, S., 2015. Working memory and executive function decline across normal aging, mild cognitive impairment, and Alzheimer’s disease. BioMed research international 2015, 748212.

Kishi, T., Hirooka, Y., Nagayama, T., Isegawa, K., Katsuki, M., Takesue, K., Sunagawa, K., 2015. Calorie restriction improves cognitive decline via up-regulation of brain-derived neurotrophic factor: tropomyosin-related kinase B in hippocampus ofobesity-induced hypertensive rats. International heart journal 56(1), 110–115.

Kuhla, A., Lange, S., Holzmann, C., Maass, F., Petersen, J., Vollmar, B., Wree, A., 2013. Lifelong caloric restriction increases working memory in mice. PloS one 8(7), e68778.

Lakowski, B., Hekimi, S., 1998. The genetics of caloric restriction in Caenorhabditis elegans. Proceedings of the National Academy of Sciences of the United States of America 95(22), 13091–13096.

Leclerc, E., Trevizol, A.P., Grigolon, R.B., Subramaniapillai, M., McIntyre, R.S., Brietzke, E., Mansur, R.B., 2020. The effect of caloric restriction on working memory in healthy non-obese adults. CNS Spectrums 25(1), 2–8.

Lennartz, R.C., 2008. The role of extramaze cues in spontaneous alternation in a plus-maze. Learning & behavior 36(2), 138–144.

Lev-Vachnish, Y., Cadury, S., Rotter-Maskowitz, A., Feldman, N., Roichman, A., Illouz, T., Varvak, A., Nicola, R., Madar, R., Okun, E., 2019. L-Lactate Promotes Adult Hippocampal Neurogenesis. Frontiers in Neuroscience 13.

Liao, C.-Y., Johnson, T.E., Nelson, J.F., 2013. Genetic variation in responses to dietary restriction--an unbiased tool for hypothesis testing. Experimental gerontology 48(10), 1025–1029.

Liao, C.-Y., Rikke, B.A., Johnson, T.E., Diaz, V., Nelson, J.F., 2010. Genetic variation in the murine lifespan response to dietary restriction: from life extension to life shortening. Aging cell 9(1), 92–95.

Lu, W., Huang, J., Sun, S., Huang, S., Gan, S., Xu, J., Yang, M., Xu, S., Jiang, X., 2015. Changes in lactate content and monocarboxylate transporter 2 expression in Aβ25-35-treated rat model of Alzheimer’s disease. Neurological Sciences 36(6), 871–876.

Lunenfeld, B., Stratton, P., 2013. The clinical consequences of an ageing world and preventive strategies. Best practice & research. Clinical obstetrics & gynaecology 27(5), 643–659.

Martin, C.K., Anton, S.D., Han, H., York-Crowe, E., Redman, L.M., Ravussin, E., Williamson, D.A., 2007. Examination of cognitive function during six months of calorie restriction: results of a randomized controlled trial. Rejuvenation research 10(2), 179–190.

Mattison, J.A., Colman, R.J., Beasley, T.M., Allison, D.B., Kemnitz, J.W., Roth, G.S., Ingram, D.K., Weindruch, R., de Cabo, R., Anderson, R.M., 2017. Caloric restriction improves health and survival of rhesus monkeys. Nature Communications 8(1), 14063.

Mitchell, S.J., Bernier, M., Mattison, J.A., Aon, M.A., Kaiser, T.A., Anson, R.M., Ikeno, Y., Anderson, R.M., Ingram, D.K., de Cabo, R., 2019. Daily Fasting Improves Health and Survival in Male Mice Independent of Diet Composition and Calories. Cell Metab 29(1), 221–228.e223.

Mitchell, S.J., Bernier, M., Mattison, J.A., Aon, M.A., Kaiser, T.A., Anson, R.M., Ikeno, Y., Anderson, R.M., Ingram, D.K., de Cabo, R., 2019. Daily Fasting Improves Health and Survival in Male Mice Independent of Diet Composition and Calories. Cell Metabolism 29(1), 221–228.e223.

Mitchell, S.J., Madrigal-Matute, J., Scheibye-Knudsen, M., Fang, E., Aon, M., González-Reyes, J.A., Cortassa, S., Kaushik, S., Gonzalez-Freire, M., Patel, B., Wahl, D., Ali, A., Calvo-Rubio, M., Burón, M.I., Guiterrez, V., Ward, T.M., Palacios, H.H., Cai, H., Frederick, D.W., Hine, C., Broeskamp, F., Habering, L., Dawson, J., Beasley, T.M., Wan, J., Ikeno, Y., Hubbard, G., Becker, K.G., Zhang, Y., Bohr, V.A., Longo, D.L., Navas, P., Ferrucci, L., Sinclair, D.A., Cohen, P., Egan, J.M., Mitchell, J.R., Baur, J.A., Allison, D.B., Anson, R.M., Villalba, J.M., Madeo, F., Cuervo, A.M., Pearson, K.J., Ingram, D.K., Bernier, M., de Cabo, R., 2016. Effects of Sex, Strain, and Energy Intake on Hallmarks of Aging in Mice. Cell Metabolism 23(6), 1093–1112.

Morgan, A.P., Fu, C.-P., Kao, C.-Y., Welsh, C.E., Didion, J.P., Yadgary, L., Hyacinth, L., Ferris, M.T., Bell, T.A., Miller, D.R., Giusti-Rodriguez, P., Nonneman, R.J., Cook, K.D., Whitmire, J.K., Gralinski, L.E., Keller, M., Attie, A.D., Churchill, G.A., Petkov, P., Sullivan, P.F., Brennan, J.R., McMillan, L., Pardo-Manuel de Villena, F., 2016. The Mouse Universal Genotyping Array: From Substrains to Subspecies. G3 Genes|Genomes|Genetics 6(2), 263–279.

Most, J., Tosti, V., Redman, L.M., Fontana, L., 2017. Calorie restriction in humans: An update. Ageing Research Reviews 39, 36–45.

Netzahualcoyotzi, C., Pellerin, L., 2020. Neuronal and astroglial monocarboxylate transporters play key but distinct roles in hippocampus-dependent learning and memory formation. Progress in Neurobiology 194, 101888.

Neuner, S.M., Heuer, S.E., Huentelman, M.J., O’Connell, K.M.S., Kaczorowski, C.C., 2019. Harnessing Genetic Complexity to Enhance Translatability of Alzheimer’s Disease Mouse Models: A Path toward Precision Medicine. Neuron 101(3), 399–411.e395.

Neuner, S.M., Wilmott, L.A., Hope, K.A., Hoffmann, B., Chong, J.A., Abramowitz, J., Birnbaumer, L., O’Connell, K.M., Tryba, A.K., Greene, A.S., Savio Chan, C., Kaczorowski, C.C., 2015. TRPC3 channels critically regulate hippocampal excitability and contextual fear memory. Behavioural Brain Research 281, 69–77.

Ouellette, A.R., Neuner, S.M., Dumitrescu, L., Anderson, L.C., Gatti, D.M., Mahoney, E.R., Bubier, J.A., Churchill, G., Peters, L., Huentelman, M.J., Herskowitz, J.H., Yang, H.-S., Smith, A.N., Reitz, C., Kunkle, B.W., White, C.C., De Jager, P.L., Schneider, J.A., Bennett, D.A., Seyfried, N.T., Chesler, E.J., Hadad, N., Hohman, T.J., Kaczorowski, C.C., 2020. Cross-Species Analyses Identify Dlgap2 as a Regulator of Age-Related Cognitive Decline and Alzheimer’s Dementia. Cell Reports 32(9), 108091.

Parikh, I., Guo, J., Chuang, K.-H., Zhong, Y., Rempe, R.G., Hoffman, J.D., Armstrong, R., Bauer, B., Hartz, A.M.S., Lin, A.-L., 2016. Caloric restriction preserves memory and reduces anxiety of aging mice with early enhancement of neurovascular functions. Aging (Albany NY) 8(11), 2814–2826.

Park, D.C., Smith, A.D., Lautenschlager, G., Earles, J.L., Frieske, D., Zwahr, M., Gaines, C.L., 1996. Mediators of long-term memory performance across the life span. Psychology and Aging 11(4), 621–637.

Pifferi, F., Terrien, J., Marchal, J., Dal-Pan, A., Djelti, F., Hardy, I., Chahory, S., Cordonnier, N., Desquilbet, L., Hurion, M., Zahariev, A., Chery, I., Zizzari, P., Perret, M., Epelbaum, J., Blanc, S., Picq, J.L., Dhenain, M., Aujard, F., 2018. Caloric restriction increases lifespan but affects brain integrity in grey mouse lemur primates. Communications biology 1, 30.

Recla, J.M., Robledo, R.F., Gatti, D.M., Bult, C.J., Churchill, G.A., Chesler, E.J., 2014. Precise genetic mapping and integrative bioinformatics in Diversity Outbred mice reveals Hydin as a novel pain gene. Mammalian Genome 25(5), 211–222.

Ridge, P.G., Mukherjee, S., Crane, P.K., Kauwe, J.S.K., Alzheimer’s Disease Genetics, C., 2013. Alzheimer’s Disease: Analyzing the Missing Heritability. PloS one 8(11), e79771.

Rogina, B., Helfand, S.L., 2004. Sir2 mediates longevity in the fly through a pathway related to calorie restriction. Proceedings of the National Academy of Sciences of the United States of America 101(45), 15998–16003.

Scott, T., Das, S., Martin, C., Stewart, T., Williamson, D., Stein, R., Bhapkar, M., Pieper, C., Rochon, J., Roberts, S., 2014. CALERIE II: the effect of 25% calorie restriction over two years on cognitive function (629.7). The FASEB Journal 28(S1), 629.627.

Sun, L.Y., Spong, A., Swindell, W.R., Fang, Y., Hill, C., Huber, J.A., Boehm, J.D., Westbrook, R., Salvatori, R., Bartke, A., 2013. Growth hormone-releasing hormone disruption extends lifespan and regulates response to caloric restriction in mice. eLife 2, e01098.

Swerdlow, R.H., 2007. Is aging part of Alzheimer’s disease, or is Alzheimer’s disease part of aging? Neurobiology of aging 28(10), 1465–1480.

Tucker-Drob, E.M., Briley, D.A., Harden, K.P., 2013. Genetic and Environmental Influences on Cognition Across Development and Context. Current Directions in Psychological Science 22(5), 349–355.

Tuttle, A.H., Philip, V.M., Chesler, E.J., Mogil, J.S., 2018. Comparing phenotypic variation between inbred and outbred mice. Nat Methods 15(12), 994–996.

van Geldorp, B., Heringa, S.M., van den Berg, E., Olde Rikkert, M.G., Biessels, G.J., Kessels, R.P., 2015. Working memory binding and episodic memory formation in aging, mild cognitive impairment, and Alzheimer’s dementia. Journal of clinical and experimental neuropsychology 37(5), 538–548.

Wahl, D., Solon-Biet, S.M., Wang, Q.P., Wali, J.A., Pulpitel, T., Clark, X., Raubenheimer, D., Senior, A.M., Sinclair, D.A., Cooney, G.J., de Cabo, R., Cogger, V.C., Simpson, S.J., Le Couteur, D.G., 2018. Comparing the Effects of Low-Protein and High-Carbohydrate Diets and Caloric Restriction on Brain Aging in Mice. Cell reports 25(8), 2234–2243.e2236.

Walker, C.K., Herskowitz, J.H., Dendritic Spines: Mediators of Cognitive Resilience in Aging and Alzheimer’s Disease. The Neuroscientist 0(0), 1073858420945964.

Watts, P., Phillips, G., Petticrew, M., Harden, A., Renton, A., 2011. The influence of environmental factors on the generalisability of public health research evidence: physical activity as a worked example. Int J Behav Nutr Phys Act 8, 128–128.

Weindruch, R., Sohal, R.S., 1997. Seminars in medicine of the Beth Israel Deaconess Medical Center. Caloric intake and aging. N Engl J Med 337(14), 986–994.

Wilkie, S.E., Mulvey, L., Sands, W.A., Marcu, D.E., Carter, R.N., Morton, N.M., Hine, C., Mitchell, J.R., Selman, C., 2020. Strain-specificity in the hydrogen sulphide signalling network following dietary restriction in recombinant inbred mice. GeroScience 42(2), 801–812.

Wimmer, M.E., Hernandez, P.J., Blackwell, J., Abel, T., 2012. Aging impairs hippocampus-dependent long-term memory for object location in mice. Neurobiology of aging 33(9), 2220–2224.

Witte, A.V., Fobker, M., Gellner, R., Knecht, S., Flöel, A., 2009. Caloric restriction improves memory in elderly humans. Proceedings of the National Academy of Sciences of the United States of America 106(4), 1255–1260.

Wu, A., Sun, X., Liu, Y., 2003. Effects of caloric restriction on cognition and behavior in developing mice. Neuroscience Letters 339(2), 166–168.

Wu, A., Sun, X., Liu, Y., 2003. Effects of caloric restriction on cognition and behavior in developing mice. Neurosci Lett 339(2), 166–168.

Yu, X., Zhang, R., Wei, C., Gao, Y., Yu, Y., Wang, L., Jiang, J., Zhang, X., Li, J., Chen, X., 2021. MCT2 overexpression promotes recovery of cognitive function by increasing mitochondrial biogenesis in a rat model of stroke. Animal Cells and Systems 25(2), 93–101.

Zajitschek, F., Zajitschek, S.R., Friberg, U., Maklakov, A.A., 2013. Interactive effects of sex, social environment, dietary restriction, and methionine on survival and reproduction in fruit flies. Age (Dordrecht, Netherlands) 35(4), 1193–1204.

