## Supplemental Figures for "Life-long Dietary Restrictions have Negligible or Damaging Effects on Late-life Cognitive Performance: A Key Role for Genetics in Outcomes"

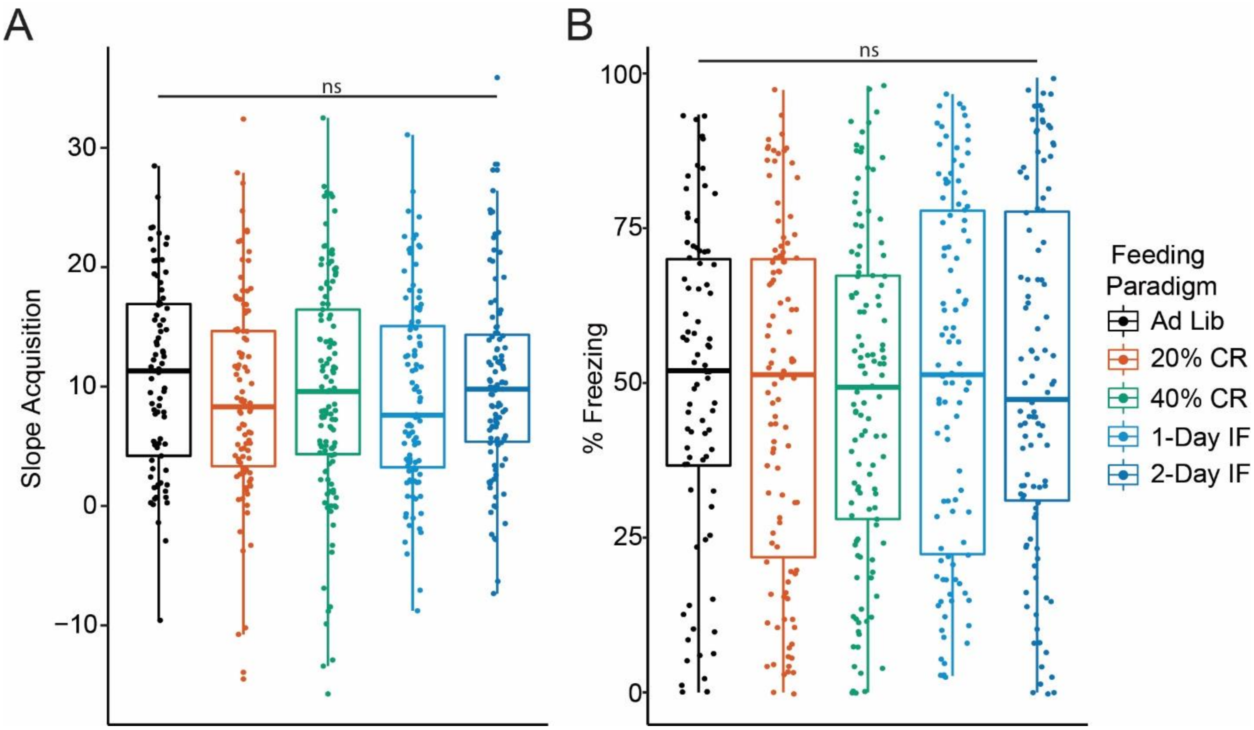


**Figure S1 | Contextual Fear Acquisition is unaltered in diversity outbred mice on caloric restriction or intermittent fasting** A) We observed no significant effect of diet on short-term slope acquisition. B) We observed no significant effect of diet on % Freezing during the post-shock 4 period. Significance was tested using linear mixed modeling with feeding paradigm as a fixed effects and test batch as a random effect. Boxes encompass the 25th to 75th percentile with whiskers indicating 10th and 90th percentiles. Median lines are indicated within each box.

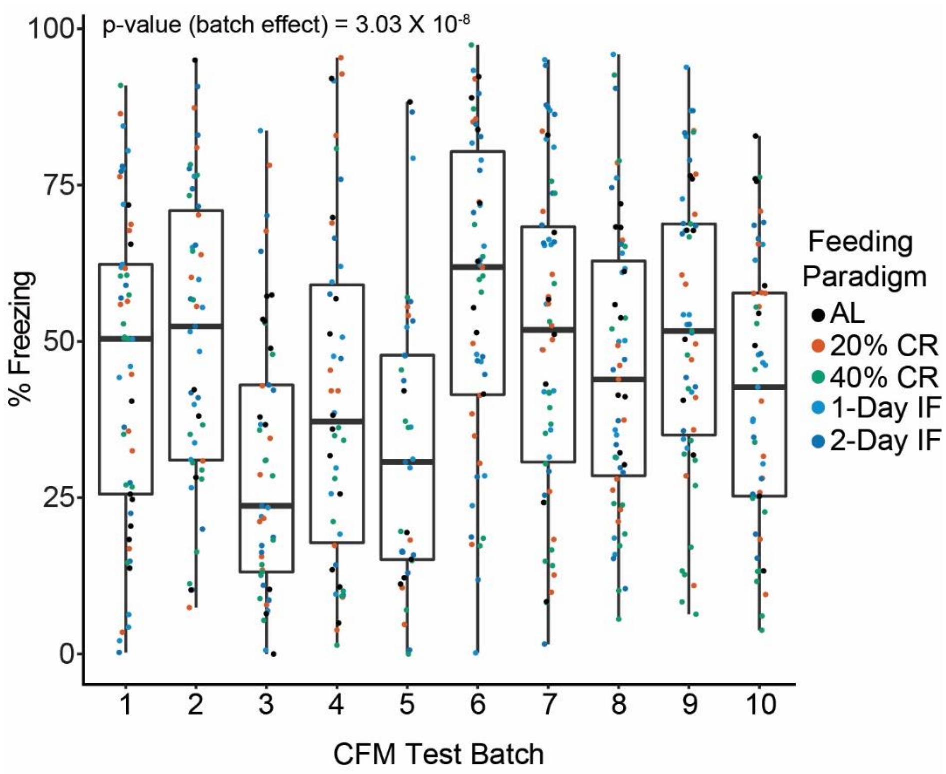


**Figure S2 | Date of CFC testing affects percent freezing.** CFM % Freezing across different days of testing. We observed significant between day variation which we account for in our statistical modeling ($F(9,492) = 6.162$, $p = 3.03 \times 10^{-8}$). Significant effect of batch was determined using 1-way ANOVA; each test day has an equally allocated proportion of diet cohorts. Boxes encompass the 25th to 75th percentile with whiskers indicating 10th and 90th percentiles. Median lines are indicated within each box
